# Pseudovirus-Mediated Proximity Labeling Identifies Candidate Host Cell Membrane Proteins Involved in Viral Attachment

**DOI:** 10.1101/2025.06.18.660084

**Authors:** Norihiro Kotani, Kensuke Iwasa, Tomoko Amimoto, Chikara Yamashita, Kumiko Komatsu, Yutaka Narimichi, Yoshiki Wakabayashi, Yuuki Kurebayashi, Yutaka Horiuchi, Reona Kashimata, Ryoko Sasaki, Megumi Kumagai, Nan Yagishita-Kyo, Takeo Awaji, Takashi Murakami, Yosuke Mizuno, Miyako Nakano, Tadanobu Takahashi, Hideyuki Takeuchi, Koichi Honke

**Affiliations:** Department of Pharmacology, Saitama Medical University, 38 Morohongo, Moroyama-machi, Iruma-gun, Saitama 350-0495, Japan; Medical Research Center, Saitama Medical University, 38 Morohongo, Moroyama-machi, Iruma-gun, Saitama 350-0495, Japan; Department of Biochemistry, Saitama Medical University, 38 Morohongo, Moroyama-machi, Iruma-gun, Saitama 350-0495, Japan; Natural Science Center for Basic Research and Development, Hiroshima University, 1-3-1 Kagamiyama Higashi-Hiroshima, Hiroshima 739-8526, Japan; Biomedical Research Center, Saitama Medical University, 38 Morohongo, Moroyama-machi, Iruma-gun, Saitama 350-0495, Japan; Department of Biochemistry, School of Pharmaceutical Sciences, University of Shizuoka, 52-1 Yada, Suruga-ku, Shizuoka 422-8526, Shizuoka, Japan; Department of Microbiology, Saitama Medical University, 38 Morohongo, Moroyama-machi, Iruma-gun, Saitama 350-0495, Japan; Graduate School of Integrated Sciences for Life, Hiroshima University, 1-3-1 Kagamiyama Higashi-Hiroshima, Hiroshima 739-8530, Japan; Department of Biochemistry, Kochi University Medical School, Nankoku, Kochi, 783-8505, Japan

**Keywords:** Proximity labeling, Membrane protein, Virus attachment, Pseudovirus, Influenza virus, SARS-CoV-2

## Abstract

Viral infections represent a significant threat to humanity, as exemplified by the COVID-19 pandemic. To mitigate the associated damage, it is essential to understand the biological characteristics of causative viruses. In this study, we report a novel method for identifying host cell molecules (including virus receptor) required for viral attachment, referred to here as “Host Cell Attachment Factors (HCAFs)”. Our approach utilizes proximity labeling (PL) technology: pseudoviruses engineered to express a PL enzyme are applied to host cells, where they bind and initiate proximity labeling. Since HCAFs are expected to be localized near viral attachment sites, candidate HCAFs are specifically tagged by the PL process. Analysis of influenza HCAFs enabled the identification of multiple candidate molecules. Subsequent virus attachment experiments using CHO cells suggested that NRP1 may serve as an HCAF for influenza viruses. Given its simplicity and rapid turnaround, this method is adaptable to a wide range of enveloped viruses, including pandemic viruses that require emergency analysis.

## Introduction

The emergence of pandemic viruses poses a significant threat to humanity, as exemplified by the COVID-19 (SARS-CoV-2) pandemic that began in 2020 and has caused extensive global damage (1, 2). Such viral crises are expected to recur in the future; for example, the emergence of novel pandemic strains of influenza viruses (family Orthomyxoviridae) remains a major concern (2–4). To mitigate the impact of these crises, it is essential to understand the biological characteristics of causative viruses. In particular, the identification and analysis of primary viral receptors and co-receptors involved in viral entry into host cells (5–8) provide crucial insights for the development of vaccines and antiviral agents.

In the case of SARS-CoV-2, angiotensin-converting enzyme 2 (ACE2) was rapidly identified as the primary receptor (7), building on previous studies of SARS-CoV-1, which caused an epidemic in the early 2000s (9). However, it is not guaranteed that future pandemic viruses will follow similar patterns. Furthermore, SARS-CoV-2 has been reported to enter cells lacking ACE2 expression, and although several candidate co-receptors (cofactors) have been proposed (10, 11), their precise roles remain unclear. Therefore, the rapid identification and characterization of host cell molecules necessary for viral attachment—referred to here as “Host Cell Attachment Factors (HCAFs)”, not limited to SARS-CoV-2, will be increasingly important for preparedness against pathogenic viruses and future pandemic viruses.

On the other hand, we have previously developed a proximity labeling (PL) technique known as EMARS (<u>E</u>nzyme-<u>M</u>ediated <u>A</u>ctivation of <u>R</u>adical <u>S</u>ources), which utilizes horseradish peroxidase (HRP)-mediated radical reactions to biochemically identify molecular complexes formed on biological membranes under physiological conditions (12, 13). This technique enables the specific labeling of molecules that are present as complexes in close proximity to a target cell membrane molecule, to which HRP has been conjugated via an antibody or similar approach. Following purification of the labeled molecules, proteomic analysis can be performed to identify the components of these molecular complexes This method is now recognized as a pioneering technology in the field of PL and has been applied to the study of molecular interactions and complexes (14–23).

Based on these advances, we hypothesized that the spike protein of a virus, attachment to the primary receptor and located in proximity to HCAFs on the host cell membrane, forms apparent molecular complexes that can be collectively labeled and identified by EMARS-based proteomic analysis. This approach could enable the comprehensive identification of candidate HCAF for SARS-CoV-2. Indeed, using this approach, we demonstrated that several membrane proteins, including DPP4, cadherin-17, and CD133, form complexes with the SARS-CoV-2 spike protein (S1-RBD) on the surface of highly susceptible Caco-2 cells, suggesting their potential involvement in SARS-CoV-2 infection (24). Thus, EMARS represents a promising tool for the identification of HCAF for various viruses, including pandemic viruses. This method allows for the rapid and straightforward listing of candidate molecules within as little as about a week, provided that the spike protein (or its binding domain recombinant protein) of the target virus is available. As such, it represents an unprecedentedly rapid and convenient approach for HCAF analysis. Moreover, since the method is not restricted by viral species as long as the spike protein is available, it is expected to be applicable to a wide range of existing viruses.

However, the current approach relies on recombinant spike proteins. The synthesis and purification of recombinant proteins requires a certain amount of time. In addition, there have been suggestions that performing EMARS under actual viral infection conditions would be more physiologically relevant and applicable to diverse viruses.

Here, we have established a system in which EMARS can be performed during the binding of not recombinant spike proteins but PL virus to host cells. PL viruses were produced using i) GPI anchor-fused HRP (14, 15)-expressing HEK293T packaging cells or ii) simultaneous transfection with the GPI anchor-fused HRP and viral genes into typical HEK293T packaging cells. PL viruses expressing the spike proteins of VSV-G, SARS-CoV-2, and influenza virus successfully bound to host cells and induced EMARS. Notably, in the analysis of the influenza virus, neuropilin-1 (NRP1) was identified as an HCAF. While sialic acid and surface glycans have long been proposed as the primary receptors for influenza virus(25, 26), our study suggests that NRP1 may serve as a novel HCAF involved in influenza virus attachment to host cells.

## Materials and Methods

### Ethics Statement

All experiments involving recombinant DNA and pseudovirus production were conducted in compliance with national regulations and the Cartagena Act. These experiments were approved by the Institutional Biosafety Committee (IBC) of Saitama Medical University (Approval No. 1647).

### Cell culture

HEK293T cells (Lenti-X™ 293T Cell Line; Takara Bio) and its transfectant cells were cultured in RPMI 1640 medium (Wako Chemicals) supplemented with 10% fetal bovine serum (FBS; GIBCO) at 37°C in a humidified 5% CO₂ atmosphere. When HEK293T cells were used as packaging cells for virus production, they should be seeded onto collagen-coated dishes (Iwaki). A549 human Caucasian lung carcinoma (JCRB0098; Japanese Collection of Research Bioresources Cell Bank, Osaka, Japan) and HeLa cells co-expressing ACE2 and TMPRSS2 (designated as HeLa-A-T; JCRB1835, Japanese Collection of Research Bioresources Cell Bank, Osaka, Japan) were maintained in RPMI 1640 medium supplemented with 5% FBS at 37°C in a humidified 5% CO₂ atmosphere. CHO-K1 cells (IFO50414; Japanese Collection of Research Bioresources Cell Bank, Osaka, Japan) were maintained in Ham’s F12 medium (Wako Chemicals) supplemented with 10% FBS at 37°C in a humidified 5% CO₂ atmosphere.

### Production of packaging cells expressing horseradish peroxidase (HRP)

The fusion genes between human codon-optimized-HRP gene and GPI anchor motif with signal sequence of DAF or Thy1 proteins were cloned into the pLenti CMV Puro DEST vector (AddGene # 17452) (14, 15). These vectors were hereinafter referred to as pLenti-DAF-HRP and pLenti-THY-HRP. HEK293T cells stably express HRP can be generated through direct transfection with pLenti-DAF-HRP and pLenti-THY-HRP using typical transfection reagents. However, conventional transient transfection may yield low HRP expression efficiency, necessitating selection and cloning. To overcome this limitation, we used lentiviral transduction for HEK293T cells to establish stable cell lines. To generate viruses carrying DAF-HRP and THY-HRP genes, 800 ng of this HRP-containing plasmid was co-transfected with 800 ng psPAX2 packaging plasmids (AddGene #12260) and 800 ng pMD2.G spike protein expression plasmid (AddGene #12259) into HEK293T cells (in 10 cm dishes and grown to ∼80% confluency) using TransIt-2020 reagent (Mirus and Takara Bio) with standard transfection protocols. The treated cells were cultured for 72 hr and then harvested supernatant containing viruses followed by filtering with 0.45 µm-filter (Sartorius). 1 mL of the filtered supernatant was added to newly plated HEK293T cells. After 72 hours of incubation, puromycin selection was initiated at 2 µg/m. Positively selected cells were maintained in puromycin-containing medium through successive passages and cryopreserved at -80°C using CultureSure® cell preservation solution (Fujifilm Wako). The engineered cells (hereinafter referred to as ”DAF-HRP packaging cells” or ”THY-HRP packaging cells”) were assessed for HRP expression through fluorescence microscopy and fluorescence-activated cell sorting (FACS) analysis (*see* details below). The cells demonstrating consistent HRP expressions were subsequently used as stable HRP-expressed HEK293T packaging cells for pseudovirus production.

### Production of PL pseudovirus

Two distinct methods were employed to generate PL viruses. In the first approach (hereinafter referred to as ”Two-step method”), the established packaging cells by stably expressing HRP in HEK293T cells described above were applied to viral production. To generate DAF-HRP or THY-HRP-expressing pseudoviruses (hereinafter referred to as ”DAF-HRP pseudovirus” or ”THY-HRP pseudovirus”) bearing the VSV-G spike protein, HRP-expressing packaging cells were transfected with pLenti CMV GFP Puro (AddGene #17448), psPAX2 (AddGene), and pMD2.G (AddGene) using standard transfection protocols. The resulting supernatants containing HRP and GFP-expressed VSV-G viruses were collected for downstream applications.

In the second approach (hereinafter referred to as ”One-step method”), to generate HRP-expressing pseudoviruses without utilizing pre-established HRP-expressing packaging cells, we directly co-transfected HEK293T cells with the following components: the pLenti-DAF-HRP plasmid constructed as described previously, psPAX2, and a spike protein expression vector of interest. For VSV-G pseudoviruses, pMD2.G was used for spike protein expression following the same procedures as described above. For the production of HRP-expressing pseudoviruses displaying the SARS-CoV-2 spike protein, the pPACK-SPIKE™ SARS-CoV-2 “S” Pseudotype Lentivector Packaging Mix (Wuhan-Hu-1; CVD19-500A-1; SBI System Biosciences) and pLenti-DAF-HRP but not psPAX2. For influenza pseudoviruses, the hemagglutinin (HA) genes from three different influenza strains, A/Puerto Rico/8/1934 (H1N1: PR8), A/Memphis/1/1971 (H3N2:M71), and A/duck/Hong Kong/313/4/1978 (H5N3: D313) were cloned respectively into the pCAGGS expression vector and used for co-transfection with the pLenti-DAF-HRP and psPAX2. In all conditions, 800 ng of each plasmid was used per 10 cm dish, and 200 ng per plasmid was used for transfection in 6-well plates. Transfections were performed using the standard protocol for TransIT-2020. Viral supernatants were harvested 48–72 hours post-transfection, filtered through 0.45 µm membranes (S7598FXOSK; Sartorius), and used for subsequent experiments. In some experiments, the pseudovirus in supernatant was purified using Lenti-X™ Maxi Purification Kit (631233; Takara Bio) according to the manufacturer’s instructions.

### Flowcytometry

Each cell was stained with i) Rhodamine-conjugate anti-HRP antibody (123-295-021; Jackson ImmunoResearch; 5 µg/ml 0.5% BSA-PBS) at room temperature for 20 min, ii) anti-ACE2 antibody (PAB886Hu01; CLOUD-CLONE, 5 µg/ml 0.5% BSA-PBS) at room temperature for 20 min followed by anti-rabbit IgG Alexa Fluor 488 (ab150077; Abcam; 10 µg/ml 0.5% BSA-PBS) at room temperature for 20 min, or iii) recombinant SARS-CoV-2 spike protein (S1-RBD; 40592-V05H; S1-RBD-mouse Fc, Sino Biological) at room temperature for 20 min followed by the secondary anti-mouse IgG Alexa Fluor 488 antibody (A-11001; Thermo Fisher Scientific; 5 µg/ml 0.5% BSA-PBS). After washing with PBS, the treated cells were analyzed using a FACS Canto II flow cytometer (BD Biosciences), to determine the expression of each molecule or binding of RGD spikes.

### Immunocytochemistry

For the confirmation of HRP expression in DAF-HRP packaging cells and THY-HRP packaging cells, stable HRP-expressing HEK293T packaging cells were detached using trypsin-EDTA (Wako), transferred to plastic tube, then stained with a goat anti-HRP primary antibody (123-005-0211; Jackson ImmunoResearch) at room temperature for 30 min followed by Alexa Fluor™ 488-conjugated donkey anti-goat IgG secondary antibody (ab150129; Abcam; 10 µg/ml 0.5% BSA-PBS). After washing with PBS, treated cells were transferred to 35-mm glass bottom dishes (627870; Greiner). HeLa-A-T cells were stained with an anti-ACE2 antibody (PAB886Hu01; CLOUD-CLONE, 5 µg/ml 0.5% BSA-PBS) at room temperature for 20 min followed by anti-rabbit IgG Alexa Fluor 488 (ab150077; Abcam; 10 µg/ml 0.5% BSA-PBS) at room temperature for 20 min, or recombinant SARS-CoV-2 spike protein (S1-RBD; 40592-V05H; S1-RBD-mouse Fc, Sino Biological) at room temperature for 20 min followed by the secondary antibody anti-mouse IgG Alexa Fluor 488 (A-11001; Thermo Fisher Scientific; 5 µg/ml 0.5% BSA-PBS). HEK293T cells directly transfected with pLenti-DAF-HRP and each virus vector were fixed with 4% paraformaldehyde (09154-85; Nacalai tesque) and then stained with Rhodamine-conjugate anti-HRP antibody (123-295-021; Jackson ImmunoResearch; 5 µg/ml 0.5% BSA-PBS) at room temperature for 30 min. These stained samples and the virus infected cells expressing GFP were observed with a fluorescent microscope BZ-700 (Keyence) or EVOS® FLoid® Cell Imaging Station (Thermo Fisher Scientific). Raw images including differential interference contrast image were captured under identical settings in the case of same experiments and then exported to TIFF or JPEG files.

### Assays of virus production and infection

In the virus production assay, the Lenti-X™ GoStix™ Plus (631281; Clontech and Takara Bio) was used to measure the amount of virus produced. A 20 μL aliquot of packaging cell culture supernatant (containing virus virion) was applied to the Lenti-X™ GoStix™ Plus stick, followed by the addition of 80 μL of the supplied buffer. The intensity of the virus-positive band was quantified using the GoStix Plus app (Clontech and Takara Bio). The amount of virus was expressed as the “GoStix value” obtained from this GoStix Plus app.

In the virus infection assay, each virus supernatant obtained from virus production method described above was filtered using 0.45 µm-filter (Sartorius) virus-containing supernatant. Lenti-X Concentrator (Takara Bio) was added to the filtered supernatant at a 1:3 (v/v) ratio and then incubated overnight at 4°C to enrich viral particles. The incubated sample was centrifuged at 1,500 × g for 45 minutes to pellet the virus. The supernatant was discarded, and the pellet was resuspended in 300 μL of serum-free medium (SFM; ASF104; Ajinomoto) with thorough mixing. The entire resuspended viral solution was added to one well of A549 cell seeded in a 12-well dish. For GFP-expressing VSV-G pseudoviruses, a 1/5 volume (60 μL) was used because of high titers. The treated cells were cultured at 37°C for 48 hours. After the infection, A549 cells were washed three times with PBS, detached using trypsin, and harvested. When assessing the infectivity of DAF-HRP pseudoviruses, we used HRP expression as the infection marker in infected host cells, as they express HRP instead of GFP. These HRP-expressing A549 host cells were stained with a rhodamine-conjugated anti-HRP antibody (123-295-021; Jackson ImmunoResearch; 5 µg/mL in 0.5%

BSA-PBS) as described above. The infected A549 host cells expressing GFP or HRP were analyzed by flow cytometry as described above. The non-virus treated A549 cells were used as negative controls. The extent of infection was quantified for convenience using FACS data as follows: the histogram images obtained from FACS analysis were processed using the image analysis software Fiji (27). For each virus-infected cell population, the area of the histogram that did not overlap with the histogram of the non-virus treated A549 cells (measured by Fiji software) was defined as infected cells. The infection rate for each virus was calculated as the ratio of the area of infected cells to the total cells (Supplementary Fig. 3).

### EMARS reaction

The EMARS reaction was performed as described previously (28, 29). Briefly, Each cell was cultured in 10 cm, 6 cm, or 6well plastic culture dishes (TPP) until approximately 80-90 % confluent, and they were subsequently treated with HRP-conjugated probes (EMARS probe) or HRP-expressing pseudovirus. For EMARS using SARS-CoV-2 spike protein, SARS-CoV-2 spike protein (0.5 µg/ml 0.5% BSA-PBS) was treated at room temperature for 30 min. After washing with PBS three times the cells were treated with HRP-conjugated anti-mouse IgG (W402B; Promega; 0.25 µg/ml 0.5% BSA-PBS) at room temperature for 30 min. After washing with PBS, the treated cells were then incubated with 0.05 mM fluorescein-conjugated tyramide (FT) (30) with 0.0075% H_2_O_2_ in PBS at room temperature for 20 min in dark.

For pseudovirus, purified and enriched pseudovirus or supernatant containing pseudovirus was added to each cell and then incubated at room temperature for 2 to 30 min (varied depending on the experiment). In the case of trypsinization for influenza viruses, 1 μl TPCK-trypsin solution (2023; Thermo Scientific; 1 mg/ml) was added to the virus in SFM solution followed by incubation at 37°C for 30 min. After treatment with these viruses, the host cells were then incubated with 0.05 mM FT with 0.0075% H_2_O_2_ in PBS at room temperature for 20 min in dark. The host cells subjected to EMARS were homogenized with 100 mM Tris-HCl (pH 7.4) through a 22 G syringe needle to rupture the plasma membrane fractions, and the samples were centrifuged at 20,000 g for 15 min to precipitate the plasma membrane fractions and used for subsequent experiments.

### Purification and enrichment of EMARS products

The precipitated cell membrane pellet was mixed with chloroform: methanol (2:1 by volume) followed by deionized water with gentle agitation. All the solvent was removed, and the resulting pellets were washed three times with 40% methanol to completely remove excess FT. To remove solution completely, the specimens were evaporated and solubilized with 100 µL of 50 mM Tris-HCl (pH 7.4) containing 1% SDS at 95°C for 5 min. The soluble material was transferred into a new tube and then diluted with 400 µL of NP-40 lysis buffer. Then, 20 µL of prepared anti-fluorescein antibody Sepharose (24) was added to the sample, which was mixed with rotation at 4°C overnight. The resins were then washed with NP-40 lysis buffer five times and 0.5M NaCl-PBS two times and used for subsequent experiments.

### Detection of EMARS products (EMARS-labeled proteins)

To confirm the presence of EMARS products, sodium dodecyl sulfate-polyacrylamide gel electrophoresis (SDS-PAGE) was performed. The precipitates containing plasma membrane fraction, or the resins (*see* “*Purification and enrichment of EMARS products* “) were resuspended by reducing SDS sample buffer (09499-14; Nacalai tesque), sonicated for approximately 5 seconds, and then heated at 100°C for 5 minutes. After cooling, each sample was resolved by 8% SDS-PAGE using Rapid Running Buffer Solution (12981-74; Nacalai tesque). The molecular weight marker was a Protein Ladder One Plus, Triple-color (19593-25; Nacalai tesque). The gel was analyzed using a ChemiDoc™ Touch (Bio-Rad; equipped with fluorescein filter) to detect fluorescein-labeled EMARS products. For loading controls, the SDS-PAGE gels after exposure were stained with Coomassie Brilliant Blue solution if necessary.

### Western blot

After SDS-PAGE gel analysis of EMARS products, the gels were subjected to western blot analysis to confirm fluorescence labeling if necessary. After electrophoresis, the gels were blotted to an Immobilon®-P PVDF Membrane (Millipore) followed by blocking with 5% skim milk solution. The membrane was incubated with sheep anti-fluorescein antibody (6400-01; SouthernBiothech; 0.5 µg /ml 5% skim milk solution) at room temperature for 1 hr followed by HRP-conjugated anti-sheep IgG (HAF016; R&D; 1:5000 5% skim milk solution) at room temperature for 1 hr.

For the detection of HRP expressions in stable HRP-expressing HEK293T packaging cells, each cell pellet collected from 10 cm dish was solubilized with 20μl reducing SDS sample buffer. Each 5 μl sample solution was subjected to 10% SDS-PAGE followed by blotting and blocking. The membrane was incubated with goat anti-HRP antibody (123-005-0211; Jackson ImmunoResearch; 1 µg /ml 5% skim milk solution) followed by HRP-conjugated anti-goat IgG (sc-2020; Santa Cruz; 1:5000 5% skim milk solution) at room temperature for 1 hr.

For the detection of HRP expressions in pseudovirus, each pseudovirus supernatant was enriched using Lenti-X Concentrator as described above (Takara Bio). The enriched virus pellet was solubilized with 50 μl reducing SDS sample buffer. Each 15 μl (VSV-G virus in Fig. S1D) or 2 μl (Influenza virus in Fig. 4A) sample solution was subjected to 8% or 10% SDS-PAGE followed by blotting and blocking. The membrane was incubated with goat anti-HRP antibody (123-005-0211; Jackson ImmunoResearch; 1 µg /ml 5% skim milk solution) followed by HRP-conjugated anti-goat IgG (sc-2020; Santa Cruz; 1:5000 5% skim milk solution) at room temperature for 1 hr.

For the detection of HA expressions in influenza pseudovirus, each pseudovirus supernatant was enriched using Lenti-X Concentrator (Takara Bio) as described above. The enriched virus pellet was solubilized with 50 μl reducing SDS sample buffer. Each 2 μl sample solution was subjected to 8% or 10% SDS-PAGE followed by blotting and blocking. The membrane was incubated with anti-influenza HA antibody mixture consisting of avian influenza A virus H5N3 HA (Hemagglutinin) antibody (GTX127299; GeneTex; 2 µg /ml 5% skim milk solution), influenza A virus H3N2 HA antibody (GTX53724; GeneTex; 2 µg /ml 5% skim milk solution), and influenza A virus H1N1 HA antibody (GTX127357; GeneTex; 0.46 µg /ml 5% skim milk solution). The incubations were performed at 4°C for overnight, After primary antibody treatment, HRP-conjugated anti-rabbit IgG (W4011; Promega; 1:5000 5% skim milk solution) was subsequently treated at room temperature for 1 hr.

Then, each membrane was developed with an Immobilon Western Chemiluminescent HRP Substrate (Millipore) and analyzed using a ChemiDoc MP Image analyzer (Bio-Rad). For loading controls, the PVDF membrane after exposure was stained with Coomassie Brilliant Blue solution if necessary.

### Transmission electron microscopy (TEM)

To investigate whether HRP expression alters viral morphology or whether HRP is detectable on the virus, each SFM cultured supernatant (ASF104; Ajinomoto) containing Inf-HRP and Inf-GFP pseudovirus was mixed 1:1 with 4% paraformaldehyde (09154-85; Nacalai tesque) at room temperature for at least 15 min. The supernatant from untransfected HEK293T cultured in SFM (serum-free medium) was used as the negative control. When labeling HRP on viral particles with gold colloid, prior to fixation, gold colloid (12 nm)-conjugated anti-HRP antibody (LS-C74279; LSBio) was added at a dilution of 1:200 and incubated at room temperature for 30 min. Subsequently, the solution was fixed with 4% paraformaldehyde as described above. A 10 μL aliquot of the fixed sample solution was placed onto a hydrophilic formvar-coated grid (Nisshin EM) and allowed to stand for 10 minutes resulting in adsorbing sample molecules onto the membrane surface. The grid was then washed by immersion in a 2% uranyl acetate solution, and the remaining solution was removed with filter paper. The grid was further washed twice in the 2% uranyl acetate solution for 1 second each. Excess stain on the grid was absorbed using filter paper, and the grid was allowed to air-dry. The prepared sample was observed at 80 kV using a JEM-1400 transmission electron microscope (JEOL).

### Proteomic analysis of EMARS products using mass spectrometry analysis

Proteomic analysis was performed using nano-liquid chromatography-electrospray ionization mass spectrometry (nano LC-ESI-MS/MS). 1% SDS solution containing MPEX PTS reagent (5010-21360; GL Science) for MS analysis was added to the washed resins (*see* “*Purification and enrichment of EMARS products* “). The samples were heated at 100°C for 5 min to elute the fluorescein-labeled molecules from resins. The eluted samples described above were treated with a final 10 mM concentration of DTT (Wako) at 50°C for 30 min, and they were then treated with 50 mM iodoacetamide (Wako) in 50 mM ammonium carbonate buffer at 37°C for 1 h. To remove SDS, SDS-eliminant (AE-1390; ATTO) was directly added to the samples, and the samples were incubated at 4°C for 1 h. After centrifugation, the supernatant was transferred to a new tube and then digested with 2 µg of trypsin (V5280; Trypsin Gold MS grade; Promega) at 37°C overnight. For peptide purification, the digested sample was applied to a GL-Tip SDB (7820-11200; GL Science) according to the manufacturer’s instructions. The sample was solubilized in 10 µL of 2% acetonitrile (ACN)/0.1% formic acid/H_2_O solution. The prepared samples were injected into an Ultimate 3000 RSLCnano (Thermo Fisher Scientific). Mass analysis was performed using a LTQ Orbitrap XL mass spectrometer equipped with a nano-ESI source (Nikkyo Technos). A NIKKYO nano HPLC capillary column (3μm C18, 75 μm I.D.×120 mm; Nikkyo Technos) was used, and this was preceded by a C18 PepMap100 column (300 μm I.D × 5mm; Thermo Fisher Scientific). The peptides were eluted from the column using a 4% to 35% acetonitrile gradient over 97 min. The eluted peptides were directly electro-sprayed into the spectrometer and were subjected to precursor scan and data-dependent MS/MS scan.

The raw data were analyzed using Proteome Discoverer ver. 3.0 software (Thermo Fisher Scientific) using MASCOT Server ver. 2.8 with the following parameters: Cys alkylation: iodoacetamide, Digestion: Trypsin, Species: Homo sapiens or Mus musculus, FDR (False discovery rate): strict 1%, relaxed 5%, respectively. The EMARS products from the experiments using three viruses were subjected to MS analysis in duplicates, and then suitable candidate HCAFs were filtered based on the following exclusion criteria: (i) keratin, immunoglobulin, histone, albumin, trypsin, and actin (ii) proteins detected in negative control samples (EMARS products from GFP-expressed influenza PR8). The classification of the identified proteins as membrane proteins was performed based on the UniProt database, and the results are listed in Tables 1 and 2.

**Table 1.** Selected candidates (TOP 20) for proximal membrane proteins around attached influenza and VSV-G pseudovirus to A549 cells.

| Accession No. | Proteins | Score |
| --- | --- | --- |
| P17301 | Integrin alpha-2 | 743 |
| O14786 | Neuropilin-1 | 524 |
| Q14126 | Desmoglein-2 | 523 |
| P17987 | T-complex protein 1 subunit alpha | 288 |
| P23229 | Integrin alpha-6 | 259 |
| P01833 | Polymeric immunoglobulin receptor | 216 |
| Q9NZU0 | Leucine-rich repeat transmembrane protein FLRT3 | 215 |
| P29317 | Ephrin type-A receptor 2 | 185 |
| P16615 | Sarcoplasmic/endoplasmic reticulum calcium ATPase 2 | 183 |
| P50895 | Basal cell adhesion molecule | 183 |
| P12830 | Cadherin-1 | 164 |
| O95490 | Adhesion G protein-coupled receptor L2 | 154 |
| P42330 | Aldo-keto reductase family 1 member C3 | 153 |
| Q92823 | Neuronal cell adhesion molecule | 149 |
| Q96QD8 | Sodium-coupled neutral amino acid symporter 2 | 140 |
| Q12864 | Cadherin-17 | 136 |
| Q9UIQ6 | Leucyl-cystinyl aminopeptidase | 131 |
| P15151 | Poliovirus receptor | 126 |
| P78310 | Coxsackievirus and adenovirus receptor | 115 |
| P15529 | Membrane cofactor protein | 108 |

\* From the search engine (Proteome Discoverer 3.0 software).

**Table 2.** Selected candidates for proximal membrane proteins around attached influenza pseudovirus to A549 cells (not around attached VSVG pseudovirus)

| Accession No. | Proteins | Score |
| --- | --- | --- |
| P15151 | CD155 (Poliovirus receptor) | 126 |
| P15529 | Membrane cofactor protein | 108 |
| O95994 | Anterior gradient protein 2 homolog | 66 |
| Q00325 | Solute carrier family 25 member 3 | 41 |
| P06753 | Tropomyosin alpha-3 chain | 33 |
| P30084 | Enoyl-CoA hydratase, mitochondrial | 14 |

\* From the search engine (Proteome Discoverer 3.0 software).

### Virus attachment assay

We used NRP1 and CD155, which were identified by proteome analysis above, as well as GPC3—a molecule not listed proteome results but previously used in SARS studies (24)— as a negative control molecule for the assay. The NRP1 (HG29858-UT; Sino Biological) and CD155 (HG10110-UT; Sino Biological) genes were purchased as inserts in the pCMV3 vector. For GPC3, we utilized pcDNA-GPC3, which was constructed in previous SARS studies (24). To generate the HRP-expressing influenza virus (PR8) for the assay, we seeded HEK293T cells in one well of a 6-well dish (TPP) and also seeded CHO-K1 cells for the assay in the remaining five wells. After 48-72 hours, the HEK293T cells were transfected with a vector set for producing pseudovirus expressing HRP and the HA of the aforementioned A/Puerto Rico/8/1934 (H1N1: PR8) strain, while the CHO-K1 cells were transfected with expression vectors for NRP1, CD155, and GPC3 (200 ng/well) using TransIt-2020 reagent (2 µl/well). After 72 hours, 20 µL influenza pseudovirus-containing culture medium was added to each well of CHO-K1 cells, and the cells were incubated at room temperature for 10 minutes to allow virus attachment. The cells were washed three times with PBS, followed by reaction with 0.05 mM biotinyl tyramide (SML2135; Sigma-Aldrich) and 0.0075% H_2_O_2_ in PBS at room temperature for 10 minutes. After three additional washes with PBS, the cells were collected into plastic tubes and treated with anti-NRP1antibody (AF3870; R&D systems; 1.25 µg/ml 0.5% BSA-PBS), anti-CD155 antibody (MAB25301; R&D systems; 1.25 µg/ml 0.5% BSA-PBS), and anti-GPC3 antibody (10088-T38; Sino Biological; 1:200 0.5% BSA-PBS) respectively at room temperature for 20 minutes. Subsequently, the cells were treated with a mixture of fluorescently labeled secondary antibodies—Alexa Fluor® 647-conjugated anti-sheep antibody (ab150179; Abcam; 5 µg/ml 0.5% BSA-PBS), anti-rabbit antibody (ab150079; Abcam; 5 µg/ml 0.5% BSA-PBS), and anti-mouse antibody (A-21235; Thermo Fisher Scientific; 5 µg/ml 0.5% BSA-PBS) —and streptavidin-Alexa Fluor® 488 (S32354; Thermo Fisher Scientific; 5 µg/ml 0.5% BSA-PBS) at room temperature for 20 minutes. After washing the cells with PBS, the fluorescence intensities of Alexa Fluor® 488 and Alexa Fluor® 647 on the cell surface were analyzed by flow cytometry (FACS) without compensation operation as described above. We compared the fluorescence intensities of Alexa Fluor® 488 (which reflects the number of biotin moieties, corresponding to the amount of HRP bound to the cells and thus the amount of virus bound to the cells) and Alexa Fluor® 647 (which reflects the expression levels of each candidate HCAF) to determine whether they were correlated. FlowJo software (ver. 10.10.0; BD Biosciences) was used in the digitization of each cell fluorescence.

### Data processing

Statistical analyses were performed using Microsoft Excel (version 2505; Microsoft Corporation) or Perplexity AI (https://www.perplexity.ai/). The data derived from virus production and infection assay were analyzed using a two-sample Student’s t-test, with the assumption of equal variances between groups in Microsoft Excel. The correlation coefficients and p-values for each sample were calculated using Pearson correlation coefficient with the assistance of Perplexity AI. Statistical significance tests for the correlation coefficients among five samples were using the Mann–Whitney U test in Perplexity AI. Statistical significance was defined as p < 0.05.

### Data availability

Proteomics raw data and search files for protein identification of HCAFs have been deposited to the ProteomeXchange Consortium (announced ID: PXD064736) via the Japan ProteOme STandard (jPOST) Repository/Database (https://jpostdb.org/) (announced ID: JPST003790).

### AI-assisted manuscript preparation

AI tool, Perplexity AI (https://www.perplexity.ai/), was employed to assist in the statistical analyses and the preparation of the manuscript, including editing, headline suggestion, and technical term generation. These tools were used as aids, and the final content was reviewed and approved by the authors.

## Results

### Production and Characterization of Packaging Cells Expressing HRP

To generate lentivirus-based HRP-expressing viruses for PL, we adopted a method in which HRP is first expressed directly in packaging cells, followed by introduction of viral vectors (Two-step method; Supplementary Fig. 1A). Initially, we constructed an HRP-expressing lentiviral vector by cloning the ORF region of human codon-optimized HRP with signal sequence and GPI-anchor region of complement decay-accelerating factor (P08174; DAF) or Thy1 (P04216; THY) into a pLenti-based plasmid. Using these vectors, we produced lentiviral particles for HRP expression. These viruses were then used to infect HEK293T cells, and stable cell lines expressing HRP were established through antibiotic (puromycin) selection. The established HRP-stable HEK293T cells were cultured, and HRP expression was confirmed (Supplementary Fig. 1B, 1C, and 1D). Fluorescence staining was observed in both DAF-HRP packaging cells and THY-HRP packaging cells (Supplementary Fig. 1B). However, stronger HRP expressions were detected in DAF-HRP packaging cells compared to THY-HRP packaging cells (Supplementary Fig. 1B and 1D). FACS analysis for DAF-HRP packaging cells revealed that at least approximately 30% of cells were highly expressed population (P3 region; Supplementary Fig. 1C). In addition, western blot analysis also showed stronger HRP expressions in DAF-HRP packaging cells (Supplementary Fig. 1D; left area).

### Production and Characterization of pseudoviruses generated by Two-step method

Next, pseudoviruses produced from both HRP-expressing packaging cell lines described above were analyzed by Western blotting to confirm HRP incorporation. HRP was detected in enriched pseudoviruses from both DAF-HRP and THY-HRP packaging cells, but pseudoviruses derived from DAF-HRP packaging cells showed modestly stronger HRP signal intensity compared to those from THY-HRP packaging cells (Supplementary Fig. 1D; right area). To examine the infectivity of pseudoviruses, VSV-G pseudoviruses for GFP expression were produced from DAF-HRP or THY-HRP packaging cells, enriched, and then applied to HEK293T cells that had been separately cultured as host cells. After 48 hours, fluorescence microscopy revealed that VSV-G pseudoviruses produced using conventional HEK293T packaging cells infected the majority of cells, resulting in GFP-positive cells (Supplementary Fig. 2A; upper column). In contrast, GFP-expressing VSV-G pseudoviruses produced from DAF-HRP or THY-HRP packaging cells exhibited a reduced number of GFP-positive cells, indicating a decrease in infection efficiency (Supplementary Fig. 2A; middle and lower column).

Using these pseudoviruses, EMARS reactions were performed. The three types of pseudoviruses prepared above were applied to HEK293T cells. Following the EMARS reaction, EMARS products were enriched and detected by electrophoresis. EMARS using pseudoviruses produced from DAF-HRP-expressing cells showed strong detection of EMARS products (Supplementary Fig. 2B; DAF-HRP). In contrast, only trace amounts of EMARS products were detected when using pseudoviruses derived from THY-HRP-expressing cells (Supplementary Fig. 2B; THY-HRP). Subsequent Western blot analysis of the post-electrophoresis gel using an anti-fluorescein antibody confirmed the presence of fluorescein-labeled proteins specifically in samples from pseudoviruses generated with DAF-HRP-expressing cells (Supplementary Fig. 2C).

### Production and Characterization of pseudoviruses generated by One-step method

The above HRP-stably expressing packaging cells exhibited weaker adhesion and slower proliferation compared to standard HEK293T cells (15), posing challenges for stable viral production. Therefore, instead of establishing HRP-stably expressing packaging cell lines, we attempted to generate HRP-expressing pseudoviruses by simultaneously co-transfecting HRP and the relevant viral genes directly into HEK293T cells (One-step method; Fig. 1A). In this case, pLenti-DAF-HRP vector has two roles: i) pLenti-DAF-HRP vector alone induces DAF-HRP expression in HEK293T cells, ii) pLenti-DAF-HRP vector simultaneously plays a role to produce pseudovirus which has DAF-HRP gene into virus genome resulting in having an ability to express DAF-HRP to infected cells. HRP expression in packaging cells following transfection was assessed by fluorescence staining. We examined three types of HEK293T cells: those co-transfected with viral vectors expressing both GFP and Influenza virus (PR8) HA as controls, and those co-transfected with DAF-HRP together with either VSV-G spike protein or influenza virus (PR8) HA viral vectors. HRP expressions were detected in both DAF-HRP-expressing packaging cell types (Fig. 1B; pLenti-Inf-HRP and pLenti-VSVG-HRP). The expression efficacy seemed to be similar to that of GFP in the control cells, which produced GFP-Influenza virus (Fig. 1B). To morphologically assess HRP expression on DAF-HRP-expressing pseudoviruses, transmission electron microscopy (TEM) was used. When comparing DAF-HRP-expressing influenza pseudoviruses and GFP-expressing influenza pseudoviruses under TEM, no morphological differences were observed (Fig. 1C). In the SFM supernatant from HEK293T cells that had not been transfected with the viral vector (used as the negative control), almost no particles of approximately 100 nm in size were observed. Furthermore, upon staining HRP on the surface of DAF-HRP-expressing influenza pseudoviruses using gold colloid-conjugated anti-HRP antibodies, virions with bound gold colloids were detected on the viral surface (Fig. 1D). To evaluate the efficiency of DAF-HRP virus production, viral production was compared using Lenti-X GoStix. The production of GFP-expressing influenza virus and HRP-expressing influenza virus was assessed, revealing a slight tendency for lower production in the HRP-expressing influenza virus compared to that of GFP (Inf-GFP vs Inf-HRP; Fig. 2A). In contrast, the production of HRP-expressing VSV-G virus was found to be significantly reduced compared to both influenza pseudovirus (Inf-HRP; Fig. 2A).

**Fig. 1.**
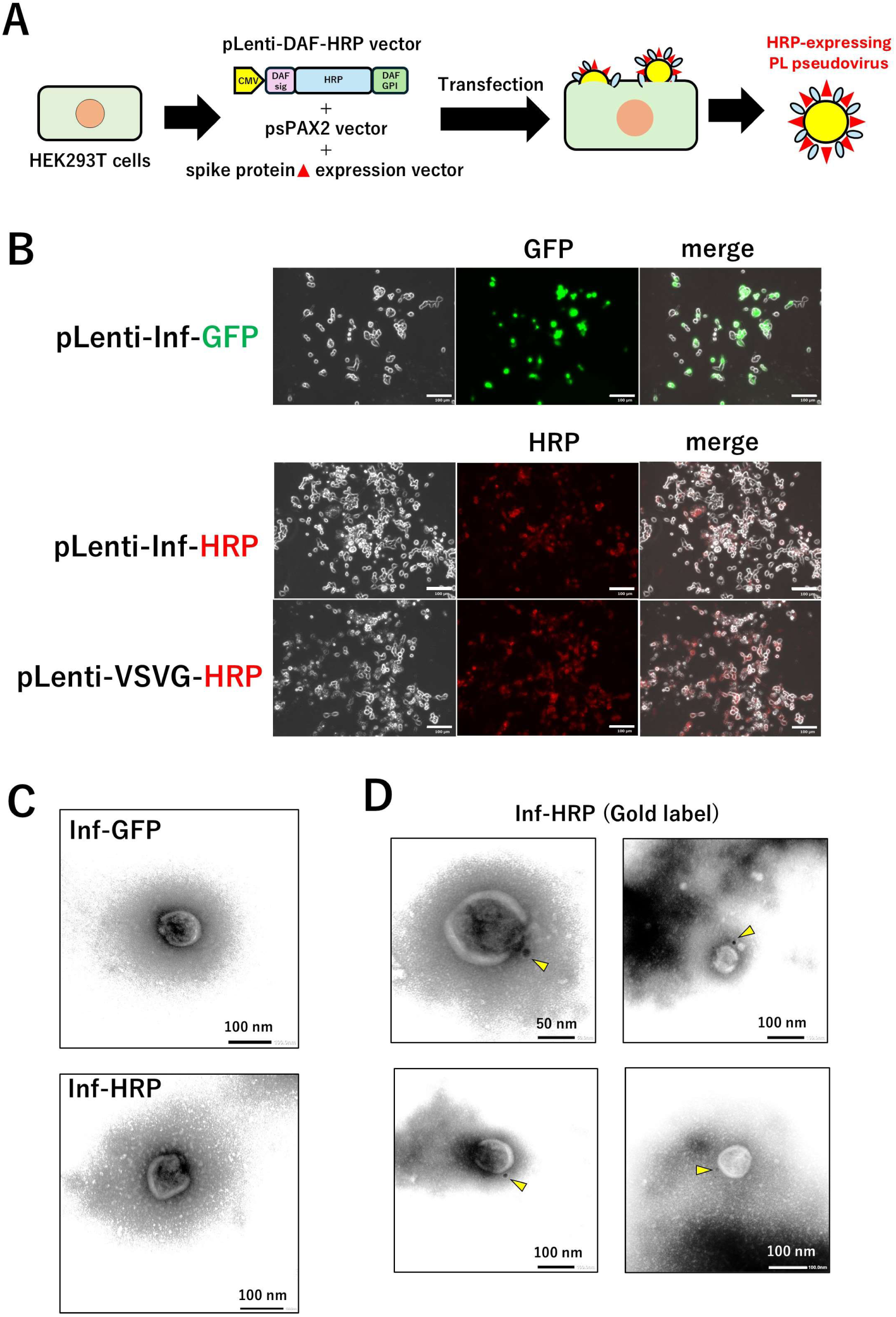
Generation of HRP-expressing pseudovirus. (*A*) Schematic illustration of the procedure for generating pseudoviruses (One-step method). For the production of HRP-expressing pseudoviruses in One-step method, HEK293T cells (a standard lentiviral packaging cell line) were co-transfected with pLenti-DAF-HRP, viral genes, and the spike protein expression vector of the virus under investigation. (*B*) Detection of HRP expressions in HEK293T cells after viral vector transduction. Following co-transfection of HEK293T cells with the above vectors, the cells were fixed after virus production and stained with an anti-HRP antibody for observation by fluorescence microscopy. Two independent experiments were performed. As a positive control, cells transfected with the pLenti-GFP and virus vector to produce pseudovirus were prepared, and GFP expression was similarly observed. The white bar indicates 100 μm. Two independent rounds of virus production and measurement were performed. (*C, D*) Morphological observation of pseudoviruses using transmission electron microscopy (TEM). The supernatant containing Inf-GFP (upper panel in (*C*)) and Inf-HRP (lower panel in (*C*)) were fixed and observed using TEM. Scale bar, 100 nm. Similarly, Inf-HRP virus produced in the same manner was treated with anti-HRP gold colloid (12 nm: yellow arrowheads) antibody and then observed by TEM. Four individual virus particles with gold colloid labeling were shown (*D*). Scale bar, 50 or 100 nm.

**Fig. 2.**
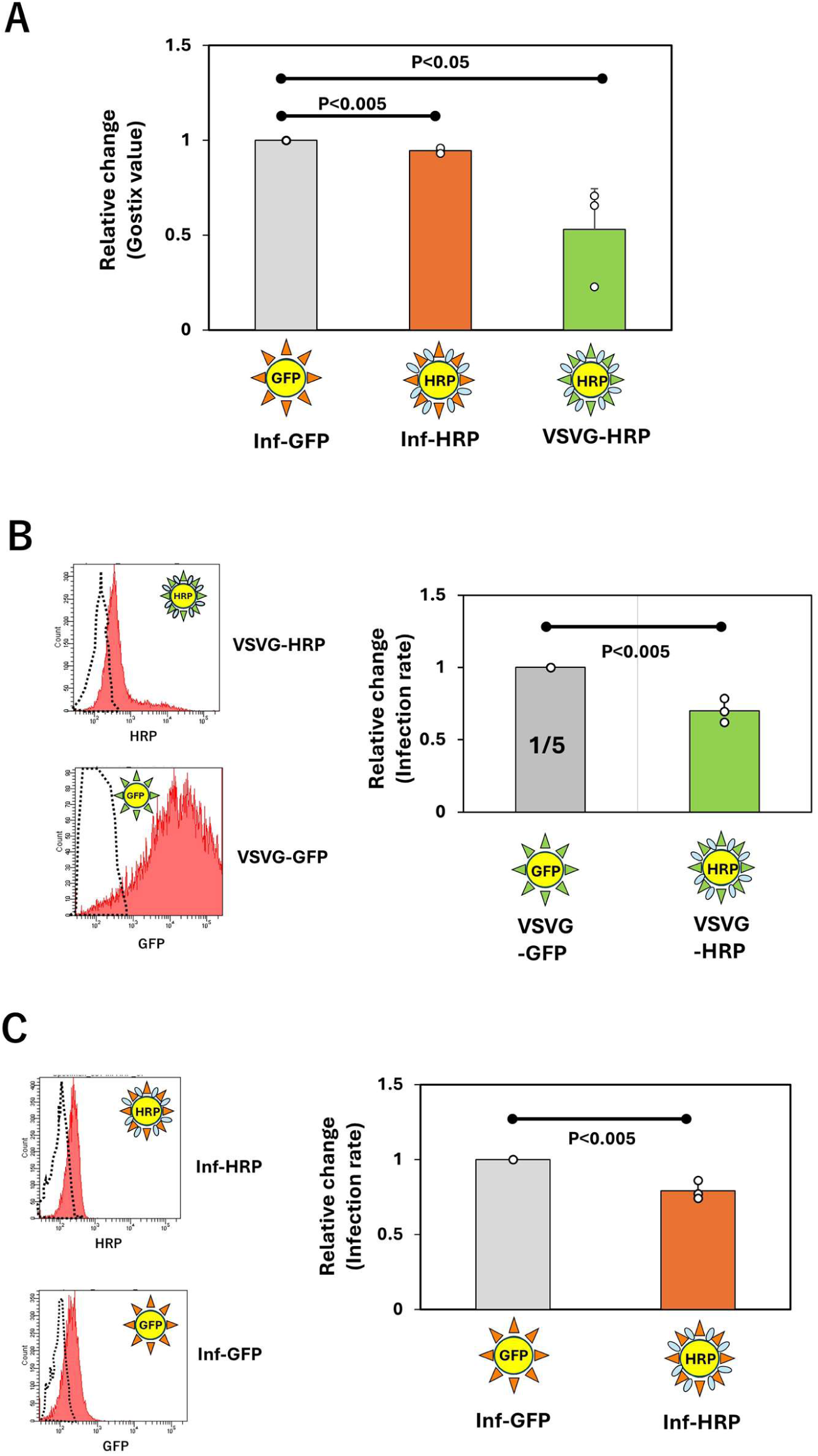
Evaluation of pseudovirus production efficiency and infectivity. (*A*) Comparison of the production efficiency of each pseudovirus using Gostix. Culture supernatants containing each virus (Inf-GFP, Inf-HRP, VSVG-HRP) were applied to Gostix as described “Materials and Methods”, and the detected positive bands were measured using the Gostix app to calculate the Gostix value. The Gostix values of HRP-expressing viruses (Inf-HRP and VSVG-HRP) were expressed as relative values using Inf-GFP as the control. For each virus, three independent rounds of virus production and measurement were performed. Statistical significance between samples was defined as P < 0.05. (*B, C*) Evaluation of infection rates for each pseudovirus. A549 cells were treated with the produced VSV-G (*B*) and influenza (*C*) pseudoviruses, and infection rates were assessed based on HRP and GFP expression levels in A549 cells post-infection (Supplementary Fig. 3). For each virus, three independent rounds of virus production and measurement were performed. The infection rates of VSVG-HRP (*B*) and Inf-HRP (*C*) viruses were expressed as relative values using VSVG-GFP and Inf-GFP as the control, respectively. Statistical significance between samples was defined as P < 0.05.

To quantitatively compare viral infectivity, we performed fluorescence-activated cell sorting (FACS) analysis. A549 cells were infected with each pseudovirus, after which HRP expression levels in cells treated with HRP-DAF-expressing pseudovirus and GFP expression levels in cells treated with GFP-expressing pseudovirus were measured as indicators of infectivity. Uninfected cells served as negative controls, and GFP or HRP expression levels in infected cells were quantified by flow cytometry. The ratio between infected cells and whole cells including uninfected cells was determined based on the analysis of histogram data, as the infection rate (Supplementary Fig. 3). When comparing HRP-expressing VSV-G with GFP-expressing VSV-G, the titer of GFP-expressing VSV-G was clearly higher, therefore, the volume of the GFP-expressing VSV-G viral solution was reduced to a volume corresponding to one fifth (considering that the production capacity of HRP-expressing VSV-G is half, the actual ratio of viral particles applied to the cells is estimated to be approximately 1:2.5). Following infection, the cells were analyzed by FACS, and as calculated in supplementary Fig. 3, it was found that the infection efficiency of HRP-expressing VSV-G was approximately 70% that of GFP-expressing VSV-G (Fig. 2B). For influenza virus, we compared HRP-expressing influenza with GFP-expressing influenza. After infection, the cells were analyzed by FACS, and as calculated in the same way as VSV-G, it was found that the infection efficiency of HRP-expressing influenza was approximately 80% that of GFP-expressing influenza (Fig. 2C).

### Pseudovirus-Mediated EMARS reaction

Using pseudoviruses produced by Two-step methods, the EMARS reaction successfully induced EMARS using pseudoviruses (Supplementary Fig. 2). Therefore, we also performed EMARS reactions with pseudoviruses produced by the one-step method. A549 cells were treated with DAF-HRP pseudoviruses carrying either VSV-G or influenza HA (PR8), and after the EMARS reaction, the EMARS products were analyzed by SDS-PAGE. The results showed that no significant EMARS products were detected when cells were not treated with any viruses or when treated with GFP-expressing pseudoviruses as negative controls (Fig. 3A). In contrast, obvious or moderate EMARS product bands were observed when cells were treated with DAF-HRP pseudoviruses carrying either influenza HA or VSV-G (Fig. 3A). In addition, western blot analysis also revealed some fluorescein-labeled proteins in the same lanes (Fig. 3B). We also performed the EMARS reaction using purified pseudoviruses from culture medium containing DAF-HRP pseudoviruses carrying influenza HA using a commercially available kit. Although the EMARS reaction was relatively weak, EMARS products were still detected (Fig. 3C). This indicates that viruses purified using such kits can be utilized for the EMARS reaction. To further clarify whether the viral particles of influenza HA-bearing DAF-HRP pseudoviruses are involved in the EMARS reaction, we modified the one-step method for virus production. Specifically, we omitted transfection of both the psPax2 packaging plasmid and the spike protein expression vector—essential components for viral packaging—and transfected only the pLenti-DAF-HRP plasmid. Under these conditions, no significant EMARS products were detected (Fig. 3D). This result confirms that functional pseudoviral particles, requiring proper packaging and spike protein incorporation, are indispensable for the EMARS reaction to occur. In addition, since efficient experimental infection by influenza virus is typically achieved by inducing HA cleavage through trypsin treatment (31), we compared EMARS products between TPCK-trypsin-treated and untreated pseudoviruses, both solubilized in serum-free medium. Given that there was no significant difference in EMARS products with or without trypsin treatment, indicating that the trypsin treatment to PL pseudovirus at least does not affect pseudovirus attachment to host cells (Fig. 3E).

**Fig. 3.**
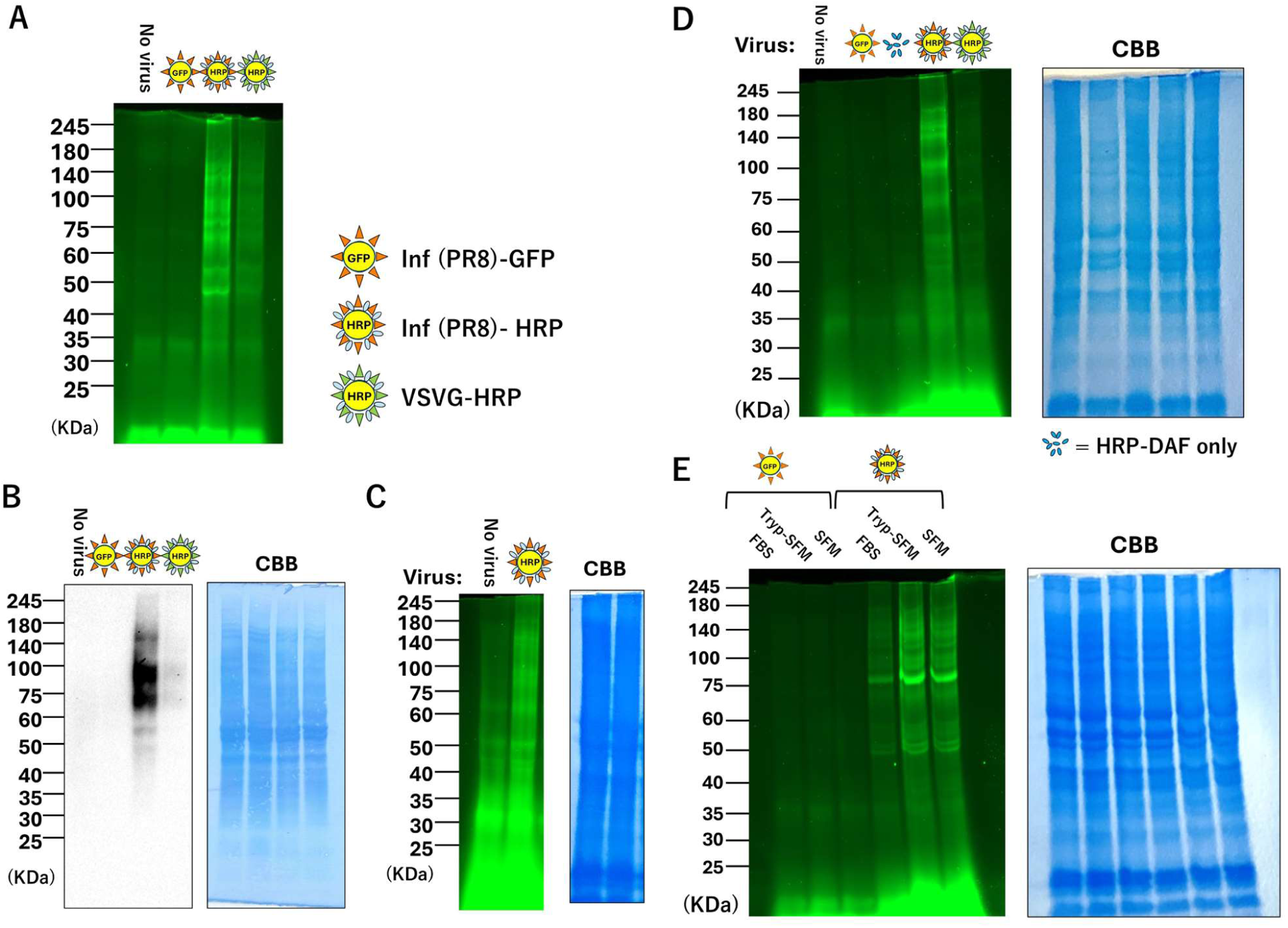
Pseudovirus-mediated EMARS reaction. (*A, B*) SDS-PAGE and western blot analysis of EMARS products. HEK293T cells seeded in the same 6-well dish were transfected with each pseudovirus vector set (Inf(PR8)-GFP, Inf(PR8)-HRP, VSVG-HRP) using the one-step method. Following culture, an equivalent amount of virus-containing culture supernatant from each virus was added to the cells. After thorough washing, FT reagent was added to initiate the EMARS reaction. The labeled molecules were then analyzed by SDS-PAGE as described “Materials and Methods” (*A*). The gel was subsequently applied to western blot analysis with anti-fluorescein antibody (*B*). Two independent experiments were performed. CBB staining was used as a loading control. (*C*) SDS-PAGE analysis of EMARS products using Inf-HRP (PR8) pseudovirus purified with a commercially available kit. The pseudovirus in supernatant was purified using Lenti-X™ Maxi Purification Kit. CBB staining was used as a loading control. Two independent experiments were performed. (*D*) SDS-PAGE analysis of EMARS products using culture supernatants of HEK293T cells transfected only pLenti-DAF-HRP. EMARS was performed using the culture supernatant of HEK293T cells that expressed only HRP-DAF via transfection with pLenti-DAF-HRP alone in which viral particle formation does not occur. This experiment was performed to determine whether non-viral components are involved in the EMARS reaction. Two independent experiments were performed. (*E*) Each influenza pseudovirus (Inf-HRP and Inf-GFP) was concentrated, resuspended in serum-free medium (*SFM*), and processed for EMARS reaction either with trypsin (*Tryp-SFM*) or without trypsin (*SFM*) treatment. Pseudovirus suspended in standard serum-containing medium (*FBS*) was also included as a control. Two independent experiments were performed.

### Comparison of EMARS using recombinant spike protein or pseudovirus

In previous studies, EMARS analysis was performed using the recombinant spike protein of SARS-CoV-2 (24). In the present study, in order to compare the results with those obtained using the newly developed pseudovirus, both the recombinant spike protein and the pseudovirus were employed in parallel to compare their EMARS products. The host cells used in this comparison were HeLa-A-T cells expressing ACE2 and TMPRSS2. We examined whether the SARS-CoV-2 spike protein bound to HeLa-A-T cells. FACS analysis revealed that the spike protein did indeed bind to these cells (Supplementary Fig. 4A; *SARS spike*). However, ACE2 expression appeared to be low (Supplementary Fig. 4A; *ACE2*), we performed additional experiments. Immunocytochemistry of ACE2 staining by anti-ACE2 antibody showed strong positiveness (Supplementary Fig. 4B; *ACE2*) compared to SARS spike (Supplementary Fig. 4B; *SARS spike*). Since the cell staining was performed after fixation, it is likely that the antibody used recognized the fixed ACE2 more efficiently, whereas ACE2 detection by FACS without fixation resulted in weaker binding signals. We performed the EMARS reaction using both recombinant spike proteins and pseudovirus and detected the EMARS products by SDS-PAGE. The results showed that the band profiles obtained from the two systems were similar; however, the quantity of EMARS products appeared to be lower with the pseudovirus compared to the recombinant spike proteins (Supplementary Fig. 4C). This observation suggests that while both methods can yield comparable patterns of protein labeling, the efficiency of the EMARS reaction may differ depending on the EMARS probe.

### Differences in EMARS products depend on viral spike proteins and influenza HA

To investigate whether differences in EMARS products arise among pseudoviruses bearing different spike proteins and influenza HA, we produced pseudoviruses carrying three types of influenza HA as well as spike proteins from SARS-CoV-2 and VSV-G using the one-step method and then performed the EMARS reaction. First, we generated pseudoviruses carrying each of the three types of influenza HA and examined the expression of viral HRP and HA proteins by Western blotting. For PR8, M71, and D313 viruses, D313 HA was expressed at higher levels compared to the other viruses (Fig. 4A; HA). In contrast, HRP expression was markedly higher in the pseudovirus carrying PR8 (Fig. 4A; HRP). Therefore, these results suggest that the PR8 pseudovirus may contain a greater amount of HRP per virion than the other viruses.

**Fig. 4.**
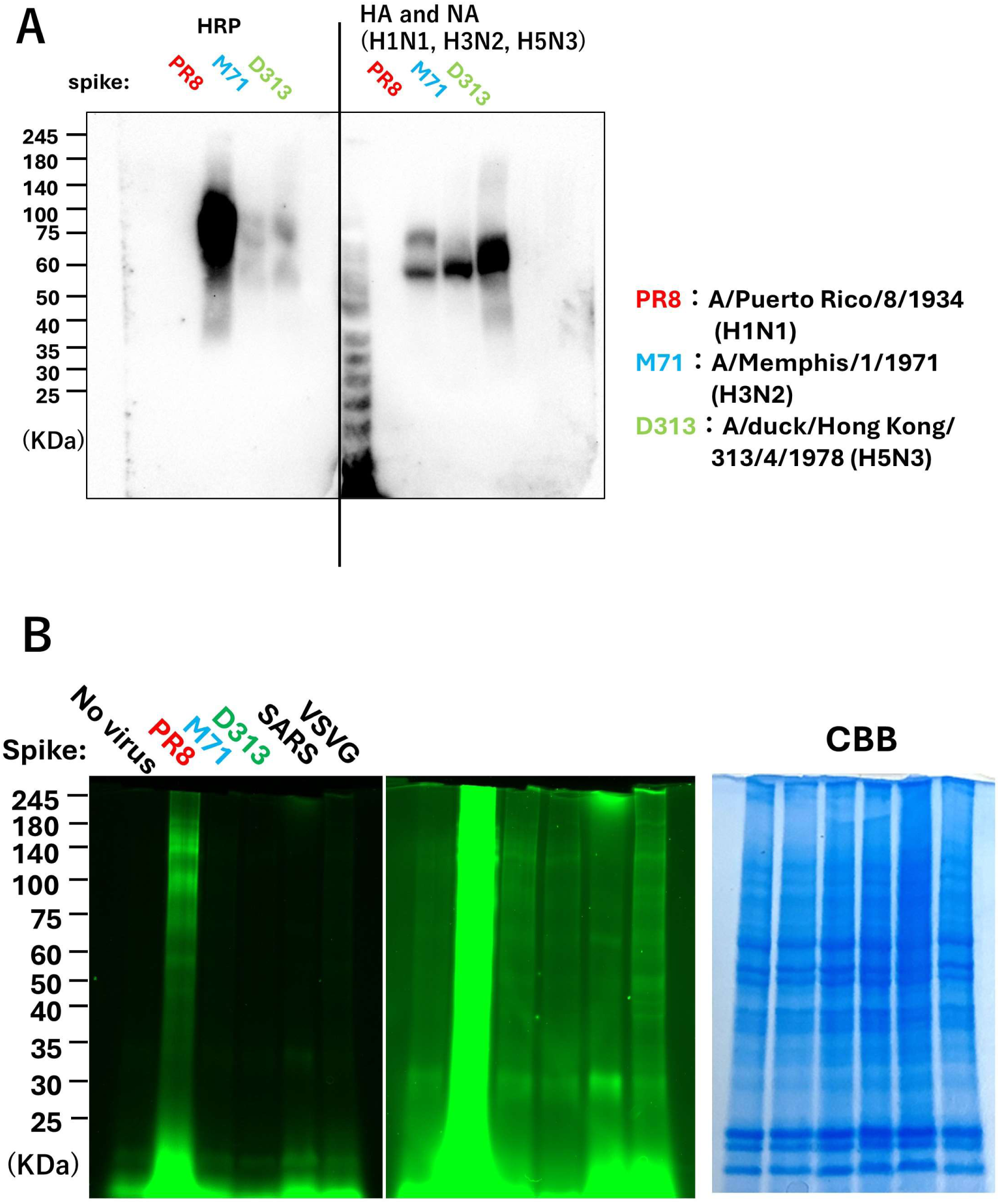
Analysis of EMARS products based on differences in HA and spike proteins. (*A*) Confirmation of HA and HRP expression in each influenza pseudovirus strain. PR8, M71, and D313 influenza pseudovirus strains were concentrated and subjected to western blot analysis. The membranes were stained using anti-HRP antibody (left panel) and anti-HA antibodies (right panel; a mixture of anti-H1N1, anti-H3N2, and anti-H5N3 antibodies) as described “*Materials and Methods*”. Two independent experiments were performed. (*B*) Detection of EMARS products using pseudoviruses carrying each HA and spike protein by EMARS. EMARS was performed using culture supernatants containing the above-mentioned PR8, M71, and D313 influenza HA pseudoviruses, as well as viruses carrying SARS-CoV-2 and VSV-G spike proteins. The EMARS products were analyzed by SDS-PAGE. The center column is an overexposed version of the data on the left column, with the detection sensitivity for samples other than PR8 increased. Two independent experiments were performed. CBB staining was used as a loading control.

In addition to these three types of influenza pseudoviruses, we treated A549 cells and HeLa-A-T cells (for SARS-CoV-2) with a total of five pseudoviruses: those carrying PR8HA, M71HA, D313HA, VSV-G, and SARS-CoV-2 spike proteins, and performed the EMARS reaction. EMARS products were then analyzed by electrophoresis. In the EMARS reaction with PR8 pseudovirus, a large number of EMARS products were detected (Fig. 4B; PR8). In the cases of M71, D313, and VSV-G pseudoviruses, fewer EMARS products were observed compared to PR8, but some bands were detected, and the band patterns appeared to differ depending on the viral species (Fig. 4B; PR8; Note: Since the host cell type for SARS-CoV-2 was different, a direct comparison may not be appropriate).

### EMARS and proteomic analysis showed the candidate HCAF

The EMARS reactions in A549 cells by three types of pseudoviruses (DAF-HRP pseudovirus carrying VSV-G and Influenza PR8 with GFP pseudovirus carrying Influenza PR8 as negative control virus) were performed and then purified and enriched EMARS products followed by subjecting proteome analysis by MS (performed in duplicate; Supplementary Table 1 to 3).

For shotgun analysis using mass spectrometry, the EMARS products were purified via immunoprecipitation with an anti-fluorescein antibody. After excluding some proteins such as keratins (see Materials and Methods), a total of 332 (first MS analysis) and 245 (second MS analysis) proteins were detected among three pseudovirus samples (Supplementary Table 1 and 2). In both VSV-G and Influenza PR8 viruses (excluding the detected proteins in EMARS products from GFP-expressed pseudovirus as negative control virus), a total of 256 (first MS analysis) and 151 (second MS analysis) proteins were detected (Supplementary Table 1 and 2). The candidate HCAFs detected only in influenza PR8 virus were a total of 161 (first MS analysis) and 45 (second MS analysis) (Supplementary Table 1 and 2). Moreover, through duplicate experiments, a total of 188 (common to VSV-G and Influenza PR8 viruses) and 6 candidate HCAFs (only in influenza PR8 virus) membrane proteins were commonly detected in both of the two proteomic analyses, respectively (Table 1 and 2).

### Virus attachment assay using the candidate HCAF-expressing CHO-K1 cells

Proteomic analysis identified candidate proteins potentially involved in viral attachment to cells. Among these, two molecules with relatively high identification scores were selected: NRP1 (common to both VSV-G virus and influenza PR8) and CD155 (detected exclusively in influenza PR8). In addition, glypican-3 (GPC3) as negative control molecule which was previously used as a candidate HCAF of SARS-CoV-2 (24). To eliminate the influence of endogenous human cell membrane proteins, we transiently expressed the three candidate molecules in Chinese Hamster Ovary (CHO) K1 cells. These cells were then treated with pseudovirus, and the attached virus was quantified. To enhance the sensitivity of viral attachment detection, we performed the EMARS reaction using biotinylated tyramide on DAF-HRP pseudovirus-treated CHO-K1 cells, with the biotin signal on the cells serving as an indicator of viral attachment levels (Fig. 5A). After performing EMARS, CHO-K1 cells were treated with primary antibodies specific to each of the three candidate molecules. The cell samples were subsequently incubated with secondary antibodies conjugated to Alexa Fluor® 647 and streptavidin-Alexa Fluor® 488. The processed cell samples were then analyzed by flow cytometry (FACS), and the fluorescence intensity of individual cells was visualized using dot plots. In this experiment, CHO-K1 cells that were not transfected with candidate molecules and were either treated or untreated with Inf-HRP were used as negative controls (Fig. 5B; “No TF and No virus” or “No TF and Inf-HRP”). Representative plots (Fig. 5B) were divided into four quadrants (vertical axis: 200, horizontal axis: 700; the “No TF and Inf-HRP” plots were treated as the “no attachment” sample) to assess viral attachment in CHO-K1 cells expressing candidate HCAFs. Since there is approximately a 0.3% signal in the second quadrant for “No TF and Inf-HRP,” the 0.3% threshold was used as the negative attachment. Among the four fractions, the fraction where both biotin and the candidate molecule were detected (double-positive in upper right area; see Fig. 5B) was operationally defined as the cell population of the candidate molecule-dependent attachment. Representative data revealed that plots in double-positive were slightly increased in CHO-K1 cells expressing NRP1 (8.4 % population) but not CD155 (4.0 % population) compared to negative control cells expressing GPC3 (6.0 % population) (Fig. 5B). However, given that these differences were minimal across all four independent experiments, we determined that the correlation between candidate HCAF expression levels and pseudovirus attachment (cell surface biotin) in individual cells would serve as the primary discriminatory index. This analysis included all cells (approximately 10,000 per experiment) subjected to FACS in each trial. Fluorescence intensities were quantified using FlowJo software, and correlation coefficients were calculated and compared with those of negative control cells (p value of correlation coefficients in all experiments were <0.05). The results based on four independent experiments indicated that only NRP1-expressing cells showed a significant positive correlation between NRP1 expression levels and viral attachment in contrast to a significant negative correlation in GPC3 as negative control molecule (Fig. 5C).

**Fig. 5.**
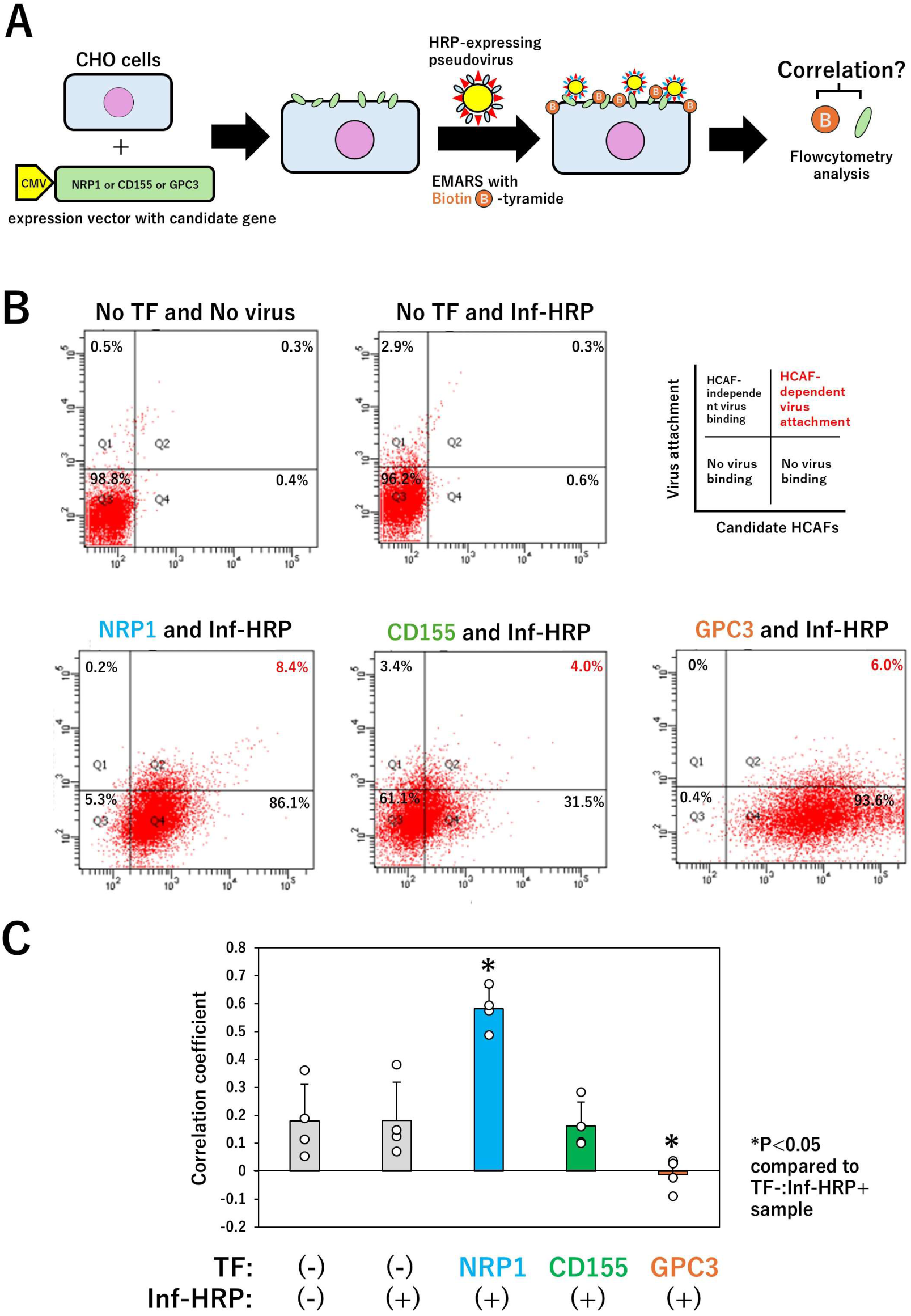
Virus attachment assays for HCAF confirmation. (*A*) Schematic illustration of the procedure for virus attachment assay. CHO-K1 cells were transiently transfected with expression vectors of NRP1, CD155, and GPC3, and then treated with Inf (PR8)-HRP pseudovirus followed by EMARS reaction with biotin tyramide as described “*Materials and Methods*”. (*B*) Representative FACS dot plot analysis. Fluorescence intensities of Alexa Fluor 488 and 647 in each cell were measured by FACS, and the results were plotted as dot plots. The vertical axis represents virus attachment (Alexa Fluor 488 fluorescence), and the horizontal axis indicates the expression level of the candidate HCAF molecule (Alexa Fluor 647 fluorescence). The plots were divided into four quadrants (vertical axis: 200, horizontal axis: 700) to assess viral attachment in CHO-K1 cells expressing candidate HCAFs. Four independent experiments were performed. (*C*) Correlation analysis between candidate molecule expression and virus attachment. The correlation between these two parameters was assessed by calculating correlation coefficients. A statistically significant correlation (p < 0.05) was observed across all samples. Furthermore, four independent analyses were performed, and the correlation coefficients from each analysis were compared. *P<0.05: compared to TF-:Inf-HRP+ sample.

## Discussion

While the impact of SARS-CoV-2 infection has become relatively stable, it remains inevitable that pandemic viruses—including novel variants such as avian influenza viruses—will continue to appear. Additionally, there are many pathogenic viruses for which HCAF is unknown or ambiguous (32). To address these challenges, it is crucial to establish an efficient system for screening HCAF candidates as an initial step.

In this study, we developed an experimental system using PL technology to address these issues. Although various PL techniques are currently available, we considered that the short interval between viral attachment and entry, as well as the relatively low number of viral particles expected to bind to host cells due to their larger size compared to proteins, would make it challenging to capture these events efficiently. Therefore, we reasoned that EMARS, a radical-based method that enables rapid reaction initiation and employs HRP to achieve high labeling efficiency, would be a suitable tool for the present study. Previous our studies (24)have relied on the use of recombinant proteins for the identification of HCAF for SARS-CoV-2. In these cases, the recombinant proteins were based on the spike protein sequence of the initially reported SARS-CoV-2 strain, and subsequent supply of spike proteins from variant strains required a certain amount of time. Therefore, for pandemic viruses with high mutation rates in the future, it is important to establish methods that can be implemented using only genetic information encoding the spike protein. Additionally, as mentioned above, there is ongoing debate regarding whether recombinant proteins truly reflect physiological virus binding, especially when compared to actual virus particles, given the significant differences in size and other properties.

We employed lentivirus-based pseudoviruses instead of recombinant spike proteins in this study. To generate viruses expressing our PL enzyme, HRP, we initially considered incorporating the HRP gene into the viral genome of the target virus. However, given the need to analyze a wide range of viruses, this approach would require significant time to individually engineer each viral genome. Moreover, depending on the size of the viral genome, inserting the HRP gene could potentially interfere with viral packaging and other critical processes. To address this, we developed a novel method for generating lentivirus-based pseudoviruses: the packaging cells were pre-engineered to anchor HRP into the viral envelope during virus generation. This approach required HRP to be expressed on the cell membrane as a soluble protein. Building on our previous research, we resolved this by constructing a fusion protein of HRP and a GPI-anchor domain, enabling stable membrane localization (14, 15). A key advantage of this system is that pseudoviruses for diverse viruses can be generated simply by modifying the spike protein expression vector, bypassing the need for time-consuming genome engineering of each target virus.

First, we established HRP-expressing HEK293T cells. HRP was fused with the signal peptide and GPI-anchoring domains from two different GPI proteins (DAF and Thy1), thereby anchoring HRP to the cell membrane. Although the underlying mechanism remains unknown, the construction using the DAF sequence resulted in more stable and robust expression (Supplementary Fig. 1). The EMARS reaction was more strongly induced with DAF-HRP pseudovirus (Supplementary Fig. 2). Therefore, the use of DAF-HRP is generally recommended. However, HEK293T cells stably expressing DAF-HRP showed reduced proliferation and poor cell viability (15, 33), making it difficult to achieve stable pseudovirus production with the Two-step method. When there is no need to incorporate a reporter gene expression sequence such as GFP into the DAF-HRP pseudovirus, more stable virus production can be achieved experimentally using the One-step method for DAF-HRP pseudovirus generation.

Next, the EMARS reactions performed using recombinant spike protein in prior studies were compared with those employing pseudovirus in the present work. Consistent with the anticipated influence of the size difference between virus particles and recombinant spike proteins on labeling efficiency, EMARS reactivity was lower with pseudovirus in the experiment (Supplementary Fig. 4C). Nevertheless, the detected banding patterns were similar (Supplementary Fig. 4C). While reduced EMARS reactivity with certain viruses may impact HCAF identification, prioritizing physiological relevance led us to conclude that initial analysis using pseudovirus-mediated assays in this study is of particular importance.

We investigated how DAF-HRP expression affects the properties and functionality of pseudoviruses produced using these methods. Our findings revealed that the impact on viral production capacity depends on the virus-specific spike protein (Fig. 2A), and thus we recommend verifying production efficiency for each virus prior to experimentation. In addition, HRP-expressing pseudoviruses can be used directly from virus-containing culture supernatants; however, they can also be conveniently concentrated using commercially available reagents when the viral titer is expected to be low or for other reasons. Regarding infection efficiency, both influenza and VSV-G pseudoviruses exhibited a 20–30% reduction compared to controls (Fig. 2B and 2C). Notably, for VSV-G pseudovirus, the observed decrease in infectivity was particularly significant when considering the lower titer of the control pseudovirus (VSV-G-GFP virus) used in the assay. These results suggest that the suitability of such pseudoviruses for HCAF labeling and identification warrants further discussion, balancing their merits and limitations in specific experimental contexts.

In contrast, for influenza virus pseudoviruses, both production efficiency and infection rates showed only a moderate decrease compared to VSV-G pseudoviruses (Fig. 2A and 2C), allowing their use for HCAF identification. In EMARS reactions, the PR8 strain influenza pseudovirus exhibited high reactivity, yielding abundant EMARS products (Fig. 3 and 4). Western blot analysis of PR8 pseudoviruses (Fig. 4A) revealed that they carry significantly more DAF-HRP on viral particles compared to other strains, though the underlying mechanism for this remains unknown. While EMARS products were also detected for VSV-G pseudoviruses, their quantity was lower than those of the PR8 strain, due to the aforementioned reductions in viral production and infectivity.

When EMARS products were compared across other viruses, including other influenza strains and SARS-CoV-2 (Fig. 4B), differences were observed, suggesting that HCAF profiles may vary depending on the virus. However, the proteomic analysis via MS identified numerous molecules common to both influenza and VSV-G pseudoviruses, particularly when the same host cells were used (Table 1). This implies that shared HCAFs may be utilized under these conditions.

Proteomic profiling yielded several candidate molecules, from which it was essential to select the HCAF(s) that significantly influence viral attachment. While the most definitive method would involve generating host cells with knockdown or knockout of the candidate molecule and subsequently assessing viral attachment, screening a large number of candidates is impractical due to time and cost considerations. This limitation is anticipated to be especially relevant in the context of analyzing pandemic viruses. Thus, as a simplified approach, this study proposes the use of CHO-K1 cells for the assay (Fig. 5A). The key features of this method are (1) the use of CHO-K1 cells to avoid interference from other human membrane proteins and (2) the application of EMARS to enhance sensitivity in detecting virus attachment. Additionally, EMARS eliminates the need for virus-specific detection antibodies (*e.g.*, anti-spike protein antibodies) to assess viral attachment.

In this analysis, when using double-positive cells (detected for both HCAF candidate expression and virus attachment via FACS) as an indicator, NRP1-expressing cells exhibited more double-positive cells compared to the negative control molecule GPC3 (Fig. 5B). However, these populations were small, suggesting that comparing correlation coefficients between HCAF candidate expression and virus attachment is a more robust analytical approach. To address this, we performed four independent experiments analyzing approximately 10,000 cells via FACS, quantified fluorescence intensities, and calculated correlation coefficients for each dataset (Fig. 5C). The computational analysis revealed a statistically significant correlation between HCAF candidate protein expression and influenza pseudovirus attachment exclusively in NRP1-expressing cells. NRP1 (Neuropilin-1) is recognized as an HCAF for several viruses, including SARS-CoV-2 (34–39). In this study, proteomic analysis identified NRP1 as an HCAF not only for influenza virus but also for VSV-G virus, suggesting its potential role as a critical molecule in the attachment and entry of viruses across a broad spectrum.

Thus, the screening and attachment assays for HCAF candidates in this study offer the advantage of being simple, cost-effective, and feasible in general biology laboratories. However, there are also several limitations. First, as exemplified by the VSV-G virus production process, the generation of HRP-expressing pseudoviruses may pose challenges depending on the virus species under investigation. Additionally, even when using the same pseudovirus and host cells, band patterns of EMARS products detected by electrophoresis can vary (Fig. 3); in such cases, repeated experiments are required for careful evaluation. Furthermore, although CD155 was identified as a candidate HCAF specific to influenza virus, no significant correlation with virus attachment was observed in this study. Therefore, the scores obtained from proteomic analysis may not always be reliable for candidate selection. It is important to emphasize that the HCAF information derived from this study should be regarded as preliminary, and final validation requires access to facilities and equipment capable of conducting infection experiments with the relevant pathogenic viruses.

In conclusion, although these limitations need to be addressed in future research, the present method is considered useful as a first screening tool for HCAF identification, given its simplicity, costs, and broad applicability. We hope that the results of this study will help to elucidate the viral attachment and entry mechanism of several envelope virus and to develop novel therapeutic agents in future.

## Acknowledgements

We thank the Saitama Medical University Biomedical Research Center for providing general technical assistance. This work was supported by Grants-in-aid for Scientific Research in Japan (No. JP21K06562 to N. K.), and Takeda Science Foundation High-Risk Emerging Infectious Diseases Research Grant.

## Author Contributions

N. K.: Conceptualization, Methodology, Validation, Resources, Data curation, Writing – review & editing, Writing – original draft, Supervision, Project administration, Funding acquisition; K. I.: Investigation, Writing – review & editing; T. A.: Investigation; C. Y.; Investigation; K. K.: Investigation; Y. N.: Investigation, Resources; Y. W.: Investigation, Resources; Y. K.: Investigation, Resources; Y. H.: Formal analysis, Resources; R. K.: Writing – review & editing; R. S.: Writing – review & editing; M. K.: Writing – review & editing; YK. N.: Writing – review & editing; T. A.: Writing – review & editing; T. M.: Formal analysis, Resources; Y. M.: Investigation, Data curation; M. N.; Investigation, Data curation; T. T.: Resources, Funding acquisition; H. T.: Resources, Funding acquisition; K. H.: Resources, Funding acquisition

---

**Fig. S1.**
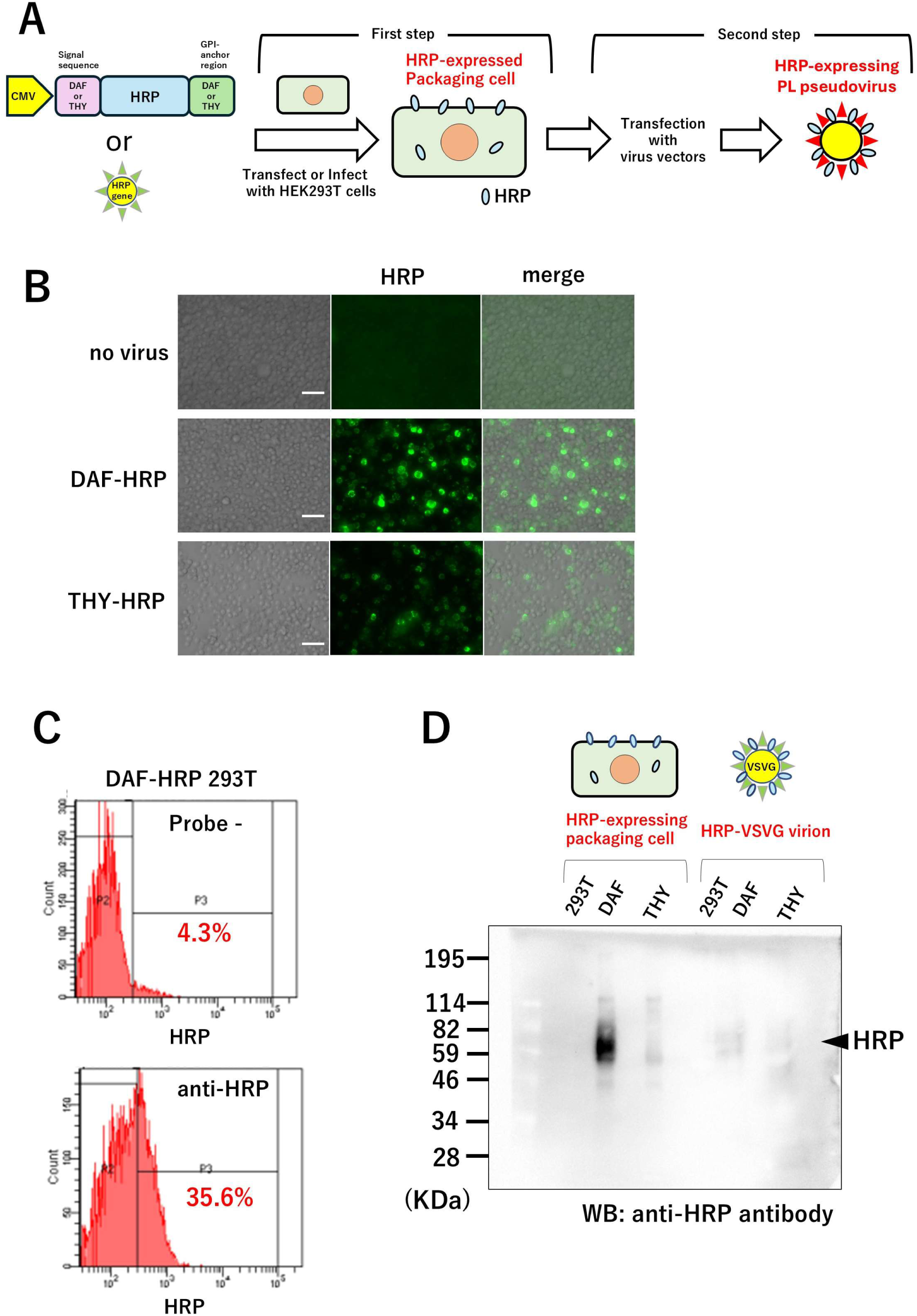
Generation of HRP-expressing pseudovirus using stable HRP-expressing HEK293T cells. (*A*) Schematic illustration of the procedure for generating pseudoviruses (Two-step method). For the production of HRP-expressing pseudoviruses in Two-step method, stable HRP-expressing HEK293T cells were first established by lentiviruses carrying DAF-HRP or THY-HJRP genes (First step). Using these stable packaging cells, the virus vectors carrying lentivirus essential gene and the spike protein expression vector were transfected for generating PL pseudovirus (Second step). (*B*) Detection of HRP expressions in lentivirus-treated HEK293T cells after 48 hr culture. Following treatment of lentiviruses carrying DAF-HRP or THY-HJRP genes with HEK293T cells, the cells were stained with an anti-HRP antibody followed by Alexa Fluor 488-conjugated second antibody for observation by fluorescence microscopy. Two independent experiments were performed. As a negative control, the cells without pseudovirus treatment were prepared (No virus). The white bar indicates 100 μm. (*C*) Representative FACS analysis for DAF-HRP expression. DAF-HRP-expressing HEK293T cells were treated with a rhodamine-conjugated anti-HRP antibody for staining. The stained cells (anti-HRP) and unstained cells served as negative control cells (Probe -) were both analyzed using FACS. The histogram data was divided into two regions, and gating was applied to calculate the percentage of cells in the high fluorescence intensity area (P3). Two independent experiments were performed. (*D*) Confirmation of HRP expression in both HRP-expressing packaging cells and PL pseudoviruses. Both DAF-HRP (*DAF*) and THY-HRP-expressing packaging cells (*THY*) were subjected to western blot analysis with anti-HRP antibody (left area). HEK293T mock cells were used as negative control cells (*293T*). DAF-HRP (*DAF*) and THY-HRP (*THY*) pseudovirus carrying VSV-G were concentrated and subjected to western blot analysis (right area). The pseudoviruses produced by typical HEK293T packaging cells were used as negative control pseudoviruses (*293T*). The membranes were stained using anti-HRP antibody as described “*Materials and Methods*”. Two independent experiments were performed.

**Fig. S2.**
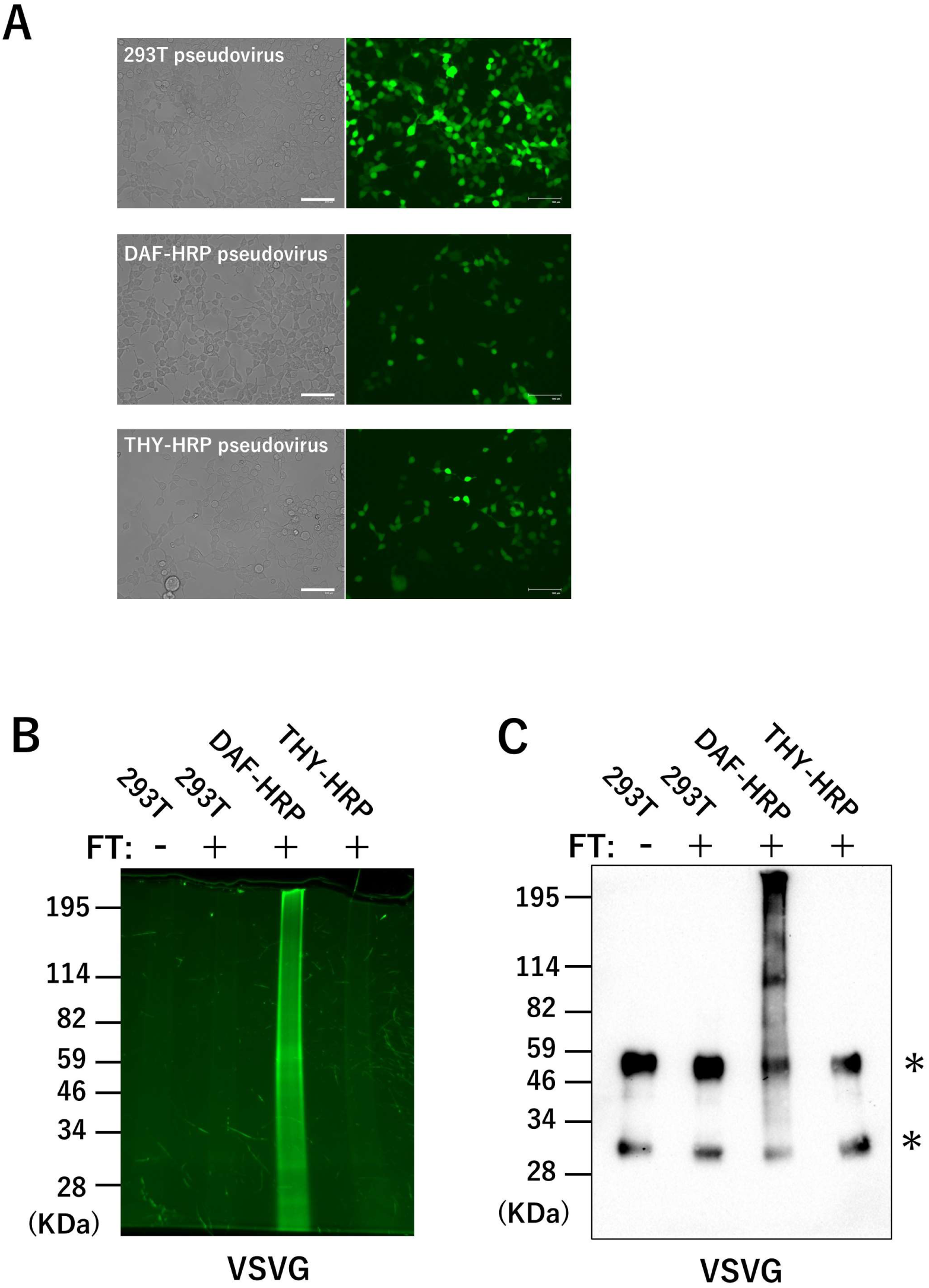
EMARS reaction using HRP-expressing pseudovirus. (*A*) Immunocytochemistry of pseudovirus-infected HEK293T cells. The VSV-G pseudoviruses carrying GFP gene derived from HEK293T cells, DAF-HRP-expressing HEK293T cells, and THY-HRP-expressing HEK293T cells were respectively infected with HEK293T cells. After 48 hr culture, each treated cell was observed by fluorescein microscopy for GFP expression. Two independent experiments were performed. The white bar indicates 100 μm. (*B, C*) SDS-PAGE and western blot analysis of EMARS products. HEK293T cells were treated with three types of VSV-G pseudoviruses in (*A*) and then performed EMARS reaction. The EMARS products were enriched with anti-fluorescein antibody-Sepharose and then analyzed by SDS-PAGE as described “*Materials and Methods*” (*B*). The gel was subsequently applied to western blot analysis with anti-fluorescein antibody (*C*). Two independent experiments were performed. Asterisks indicate the bands of H and L chain from anti-fluorescein antibody-Sepharose.

**Fig. S3.**
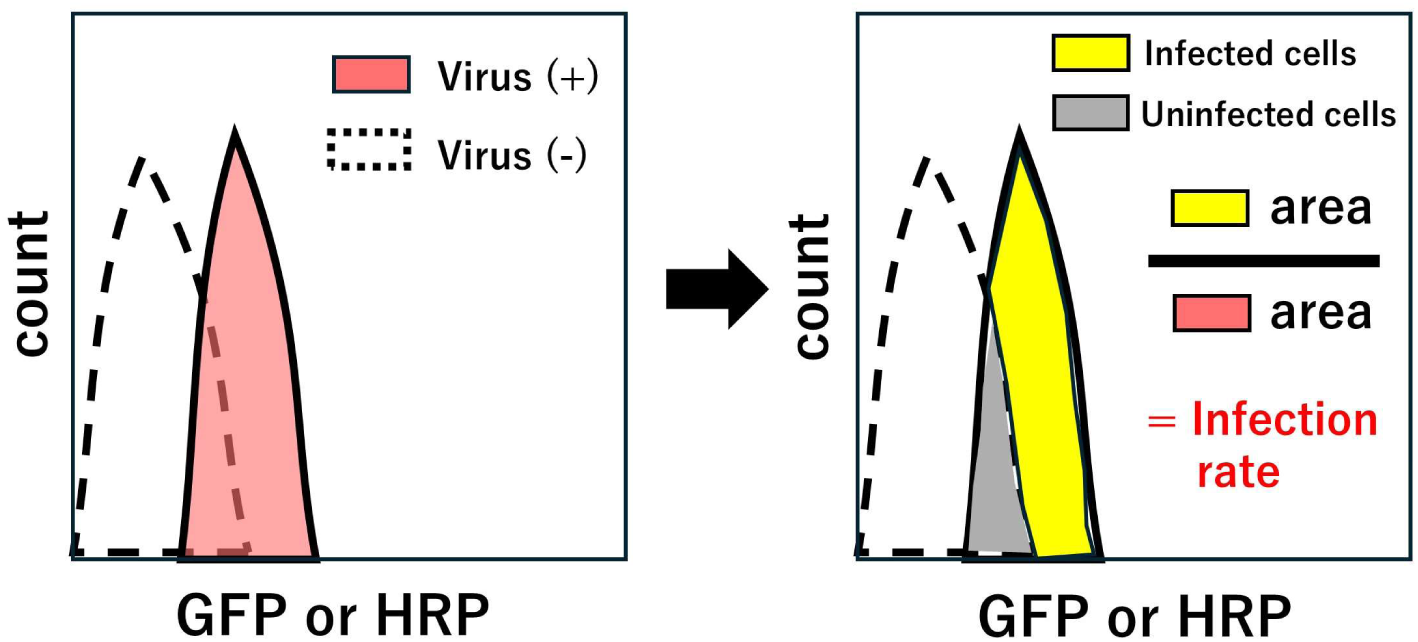
Calculation of infection rate using FACS analysis data. Histograms were imported into Fiji (NIH ImageJ), and the areas corresponding to infected and uninfected cells were quantified. The infection rate was calculated based on these area measurements.

**Fig. S4.**
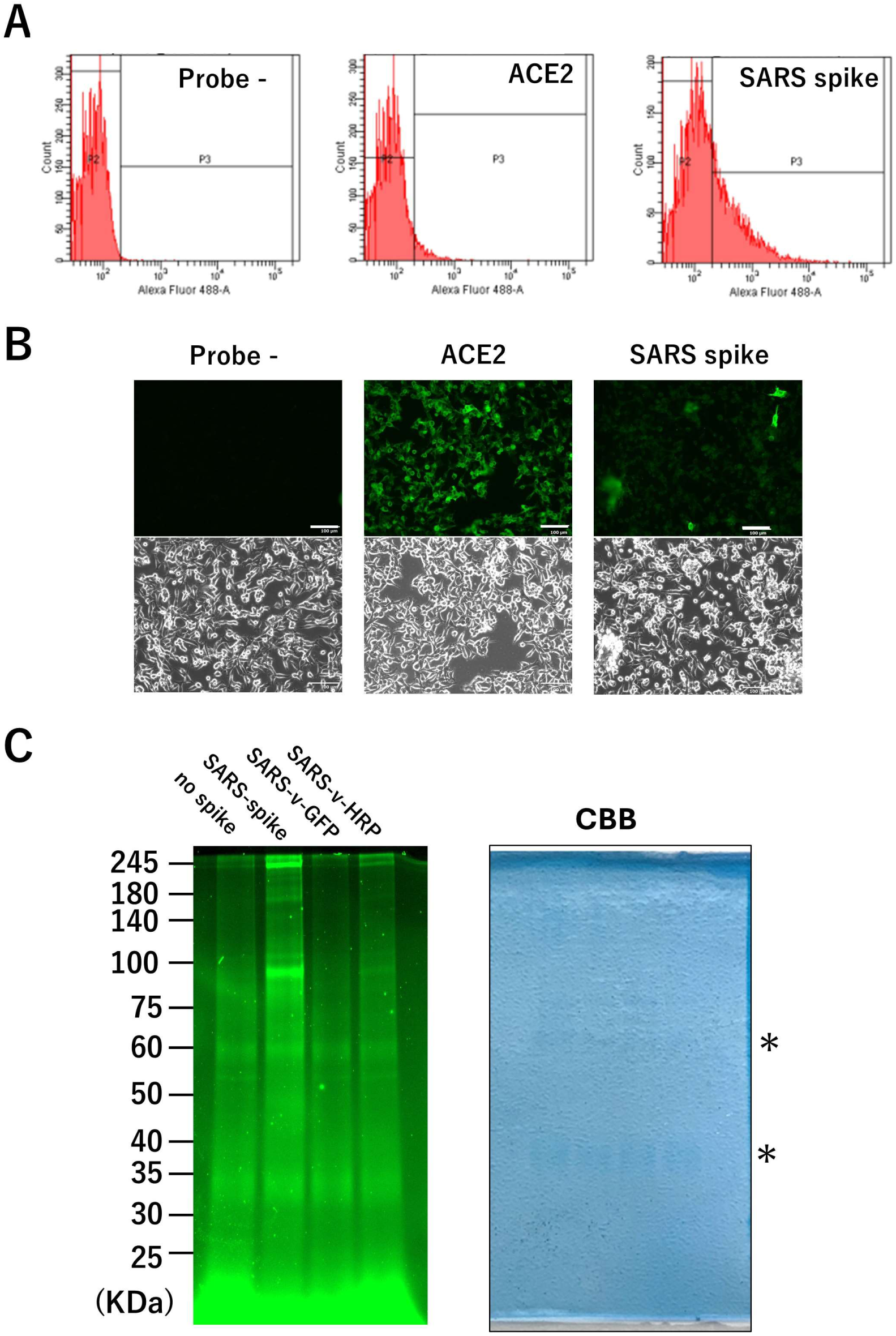
Differences between EMARS using recombinant spike protein or PL pseudovirus. (*A, B*) Characterization of HeLa-A-T cells as host cells. HeLa-A-T cells were assessed for ACE2 expression (*ACE2*) and spike protein binding (*SARS spike*) using FACS (*A*) and immunocytochemistry (*B*). Cells without antibody and spike protein treatment were served as negative controls (*Probe–*). The white bar indicates 100 μm. (*C*) SDS-PAGE analysis of EMARS products derived from both spike and pseudovirus. HEK293T cells were treated with recombinant spike protein (*SARS-spike*), HRP-expressing pseudovirus carrying SARS-CoV-2 spike proteins (*SARS-v-HRP*), or GFP pseudovirus carrying SARS-CoV-2 spike proteins (*SARS-v-GFP*) as negative control viruses. After thorough washing, FT reagent was added to initiate the EMARS reaction. The EMARS products were enriched with anti-fluorescein antibody-Sepharose and then analyzed by SDS-PAGE as described “*Materials and Methods*”. Two independent experiments were performed. CBB staining was used as a loading control; however, due to the purification of EMARS products, no protein bands detectable by CBB were observed except for nearly equal amounts of heavy and light chains (asterisks; derived from antibodies bound to the resin during the enrichment process).

